# Modeling RNA-binding protein specificity *in vivo* by precisely registering protein-RNA crosslink sites

**DOI:** 10.1101/428615

**Authors:** Huijuan Feng, Suying Bao, Sebastien M. Weyn-Vanhentenryck, Aziz Khan, Justin Wong, Ankeeta Shah, Elise D. Flynn, Chaolin Zhang

## Abstract

RNA-binding proteins (RBPs) regulate post-transcriptional gene expression by recognizing short and degenerate sequence elements in their target transcripts. Despite the expanding list of RBPs with *in vivo* binding sites mapped genomewide using crosslinking and immunoprecipitation (CLIP), defining precise RBP binding specificity remains challenging. We previously demonstrated that the exact protein-RNA crosslink sites can be mapped using CLIP data at single-nucleotide resolution and observed that crosslinking frequently occurs at specific positions in RBP motifs. Here we have developed a computational method, named mCross, to jointly model RBP binding specificity while precisely registering the crosslinking position in motif sites. We applied mCross to 112 RBPs using ENCODE eCLIP data and validated the reliability of the resulting motifs by genome-wide analysis of allelic binding sites also detected by CLIP. We found that the prototypical SR protein SRSF1 recognizes GGA clusters to regulate splicing in a much larger repertoire of transcripts than previously appreciated.

## Introduction

RNA-binding proteins (RBPs) are central for post-transcriptional regulation of gene expression by interacting with specific sequence or structural elements embedded in their target transcripts (Licatalosi and Darnell, 2010). Precise characterization of RBP binding specificity is crucial to identify protein-RNA interaction sites important for gene expression regulation, and understand how such interactions are affected by genetic variation, particularly in the context of human disease(Lunde et al., 2007). The latter is underscored by the observation that 90% of human disease- or trait-associated single nucleotide polymorphisms (SNPs) identified by genome-wide association studies (GWAS) are located in the noncoding regions of the genome (Hindorff et al., 2009), including exons and introns, and are enriched in expression or splicing quantitative trait loci (eQTLs or sQTLs) (Li et al., 2016). However, mechanistic insights into whether and how these GWAS SNPs directly affect gene expression and splicing are currently limited.

Multiple approaches have been used to determine RBP binding sites and define RBP binding specificity. *In vitro* RNA selection is an iterative procedure to purify sequences with high-affinity binding to an RBP of interest, starting from a large library of random oligos (Wilson and Szostak, 1999). Recently, several high-throughput assays, such as RNAcompete (Ray et al., 2009; Ray et al., 2013), RNA Bind-and-Seq (RBNS) (Dominguez et al., 2018; Lambert et al., 2014), and RNA-MaP (Buenrostro et al., 2014), were also developed to define RBP specificity *in vitro* and these assays have been applied to hundreds of RBPs.

UV cross-linking and immunopreciptiation (CLIP) of protein-RNA complexes, followed by high-throughput sequencing of isolated RNA fragments (HITS-CLIP), is a biochemical assay to map *in vivo* protein-RNA interactions on a genome-wide scale (Ule et al., 2003, Ule et al., 2005, Licatalosi et al., 2008). Since its initial development, CLIP and multiple variant protocols have been applied to an expanding list of RBPs in various species and cellular contexts (Darnell, 2010, Licatalosi and Darnell, 2010). In particular, a modified version of CLIP, named eCLIP, was adopted by the Encyclopedia of DNA Elements (ENCODE) consortium to map the binding sites of over 100 RBPs in two human cell lines, HepG2 and K562, making it the largest CLIP dataset generated thus far (Van Nostrand et al., 2017; Van Nostrand et al., 2016).

Both *in vitro* binding assays (such as RNAcompete) and CLIP generate a list of sequences expected to be bound by an RBP. A common pattern shared by these sequences, or motif, needs to be inferred *de novo* by statistical modeling to define the sequence specificity of the RBP and predict novel binding sites. A similar task is present for studies of DNA-binding proteins that regulate transcription, which was historically the initial focus of genomic analysis using large scale datasets. Therefore, current methods used for *de novo* RBP motif discovery (e.g., MEME and HOMER) were originally developed for analysis of DNA-binding proteins (Bailey and Elkan, 1994; Heinz et al., 2010). However, there exist important differences between DNA-binding proteins and RBPs.

As compared to DNA-binding proteins, most RBPs recognize very short (~3–7 nt) and degenerate sequence motifs, which in general have limited information content (Chen and Manley, 2009; Lunde et al., 2007; Singh and Valcarcel, 2005). For example, the high-affinity binding motif of the neuron-specific splicing factor Nova is the tetramer YCAY (Y=C/U) (Jensen et al., 2000). Other examples include recognition of YGCY elements by Mbnl (Du et al., 2010; Goers et al., 2010), UCUY by Ptbp1 (Perez et al., 1997) and Ptbp2 (Licatalosi et al., 2012), and U-tracts by Hu (Gao et al., 1994; Levine et al., 1993) (reviewed by (Chen and Manley, 2009)). Due to the apparently lower specificity of RBPs, the performance of the current computational tools for *de novo* motif discovery varies when applied to RBPs. Consequently, despite the availability of CLIP or high-throughput *in vitro* binding data, the specificity of many RBPs remains to be defined. This challenge is reflected in situations in which distinct motifs have been reported for the same RBPs from different datasets (e.g., for FMRP (Ascano et al., 2012; Darnell et al., 2005; Darnell et al., 2001; Darnell et al., 2011)). In addition, multiple RBPs were reported to have similar motifs and yet they have very distinct binding maps in the transcriptome (e.g., for TIA1, hnRNP C and other RBPs recognizing U-rich or AU-rich elements (Konig et al., 2010; Wang et al., 2010)).

The degeneracy of RBP binding motifs argues for the importance of mapping RBP-binding sites with high resolution to improve the accuracy of motif discovery. Previously, we developed computational approaches to map the extract protein-RNA crosslink sites through analysis of crosslink-induced mutation sites (CIMS) or truncation sites (CITS) using CLIP data (Weyn-Vanhentenryck et al., 2014; Zhang and Darnell, 2011). CIMS and CITS are signatures of protein-RNA crosslinking introduced by interference of reverse transcription by the covalently linked amino acid-RNA adducts, and they provide a means of mapping protein-RNA interactions at single-nucleotide resolution. Furthermore, our previous analysis revealed that UV crosslinking frequently occurs at specific positions in the RBP binding motifs, most likely reflecting critical RNA residuals for direct protein-RNA contacts (e.g., G2 and G6 in UGCAUG that is recognized by RBFOX) (Moore et al., 2014; Weyn-Vanhentenryck et al., 2014). Here we report that these crosslink sites can be used to precisely register RBP binding sites, at single-nucleotide resolution, to improve the accuracy of *de novo* RBP motif discovery. We demonstrate the effectiveness of this strategy by developing a statistical model and algorithm named mCross and applying it to 112 RBPs using ENCODE eCLIP data. The reliability of the resulting motifs defined by mCross was validated by analysis of allelic protein-RNA interaction sites caused by heterozygous SNPs on a genome-wide scale. Based on motifs defined by mCross, we unexpectedly found that SRSF1, a prototypical SR protein, predominantly recognizes clusters of GGA half sites, instead of the canonical GGAGGA motif, which allows it to regulate splicing of a much larger repertoire of transcripts than previously appreciated. Finally, we have developed a searchable, interactive web interface (http://zhanglab.c2b2.columbia.edu/index.php/MCross) to allow the research community to have easy access to this resource.

## Results

### Joint modeling of RBP binding specificity and precise crosslinking positions

Most of the *de novo* motif discovery tools currently available use a standard model of a position-specific weight matrix (PWM) (Stormo, 2000) to characterize the specificity of DNA- or RNA-binding proteins (note that the consensus can be viewed as a special case of a PWM). mCross takes advantage of the precise protein-RNA crosslink sites inferred from CIMS and CITS analysis and the observation that crosslinking frequently occurs at specific positions within the motif (Figure 1A, B). Therefore, it augments the standard PWM model by jointly modeling RBP sequence specificity and the precise protein-RNA crosslink sites that help register motif sites in longer input sequences at single-nucleotide resolution (Figure 1C). This model allows us to dramatically limit the search space. Optimal model parameters are determined by maximizing the likelihood function (see STAR Methods).

**Figure 1:**
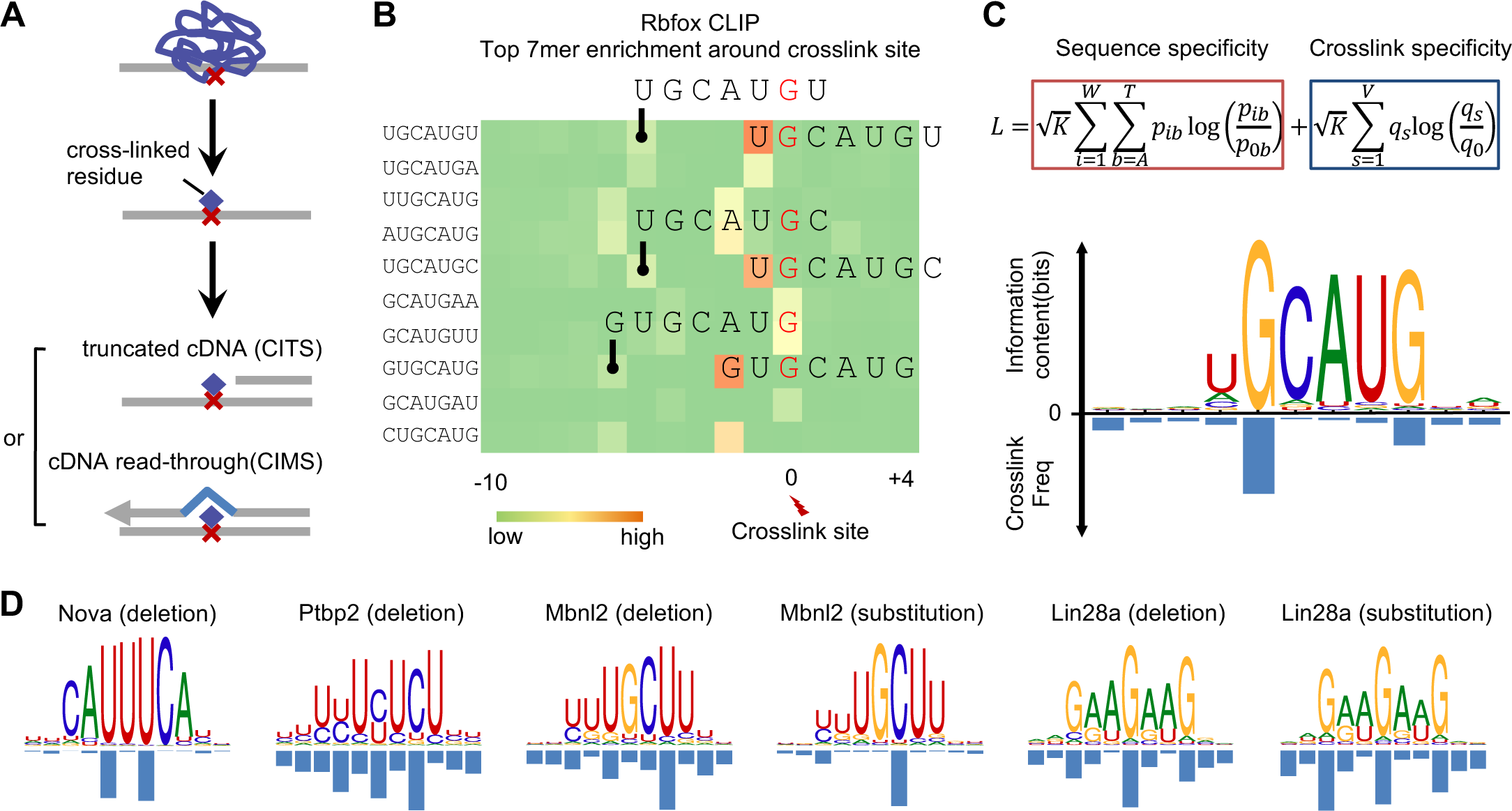
Overview of mCross to jointly model RBP binding specificity and frequency of crosslink sites at different motif positions. Related to Figures S1 and S2. (A) Schematic of identifying reproducible protein-RNA crosslink sites by analysis of crosslinking-induced mutation sites (CIMS) and truncation sites (CITS). (B) Enrichment of top 7-mers in sequences around Rbfox crosslink sites. Crosslink sites were identified by CIMS analysis of deletions, and the frequency of each 7-mer starting at different positions relative to the crosslink site is shown in the heatmap. Representative 7-mers showing the highest position-specific enrichment are indicated and the corresponding crosslinked nucleotide is highlighted in red. (C) The likelihood function that jointly models RBP sequence specificity and crosslinking positions are shown at the top. The Rbfox binding motif and the crosslinking probability in each position of the motif identified by mCross *de novo* are shown at the bottom. (D) Additional examples of RBP motifs and crosslinking positions as discovered by mCross.

For initial assessment on the reliability of mCross, we applied it to several tissue-specific RBPs, including Rbfox (Weyn-Vanhentenryck et al., 2014), Nova(Zhang et al., 2010), Ptbp2 (Licatalosi et al., 2012), Mbnl2 (Charizanis et al., 2012), and Lin28a(Cho et al., 2012), whose binding specificity varies in a wide range but has been thoroughly characterized using CLIP and other experimental approaches. In each case, mCross recovered the well-defined motif as well as the predominant crosslink sites within the motif simultaneously (Figure 1C, D). For example, Rbfox binds the (U)GCAUG element with a certain level of degeneracy at the first position, U1, and the predominant crosslink sites are G2 and G6. Ptbp2 binds to UCUCU-like elements with predominant crosslink sites at the cytosines. Importantly, all motifs of the same RBP, as discovered by mCross, are highly similar to each other, minimizing the ambiguity in determining the *bona fide* specificity (Figure S1). These results also confirmed that photocrosslinking can occur at different nucleotides, although a uridine-bias was assumed in general.

To further evaluate the effectiveness of mCross, we applied it to Argonaute (Ago) CLIP data of the mouse brain (Chi et al., 2009) to recover microRNA binding sites that are reverse complementary to the seed sequences. Since Ago mRNA CLIP tags could capture binding sites of all microRNAs expressed in the brain, the data represent a mixture of multiple motifs. mCross successfully identified motifs that can be grouped into 10 clusters, including canonical seed matches of six miRNAs that are abundantly expressed in the brain and the miR-124 seed matches with a bulge (Figure S2) (Chi et al., 2012). In general, Ago preferably crosslinks to target mRNA transcripts at positions flanking seed matches, but in some cases it appears that uridines inside the seed matches are also prone to crosslinking. This example suggests that mCross is capable of deconvoluting different modes of binding in more complex datasets.

### Defining motifs of 112 unique RBPs using ENCODE eCLIP data

Having demonstrated the promise of mCross, we extended our analysis to eCLIP data from ENCODE (Van Nostrand et al., 2016). These include 70 RBPs in HepG2 cells, 89 RBPs in K562 cells, and 1 RBP in adrenal gland, with each RBP assayed by two replicate experiments (in total, 160 experiments×2=320 independent CLIP libraries, representing 112 unique RBPs; as of Dec 30, 2016). The replicates allow for evaluation of reproducibility (see below). All CLIP data were processed using our established CTK package (Shah et al., 2017) to call CLIP tag cluster peaks and infer potential crosslink sites using CIMS and CITS analysis (Figure S3 and Table S1).

Since the majority of RBPs assayed by eCLIP were previously poorly characterized, we developed quantitative metrics to evaluate if an RBP likely has robust binding specificity. First, we estimated the number of 7-mers that are asymmetrically enriched in CLIP tag clusters, as compared to the number of 7-mers that are depleted. Second, we developed a ‘disconcordance’ score (D-score) to measure whether top 7-mers are consistently enriched in the two biological replicates, with a low D-score indicating high concordance between the replicates (see STAR Methods). For example, RBFOX2 showed a very low D-score between the two replicates (D<0.00067), and most of the significantly enriched 7-mers contained the (U)GCAUG motif that is known to bind the protein (Figure 2A).

**Figure 2:**
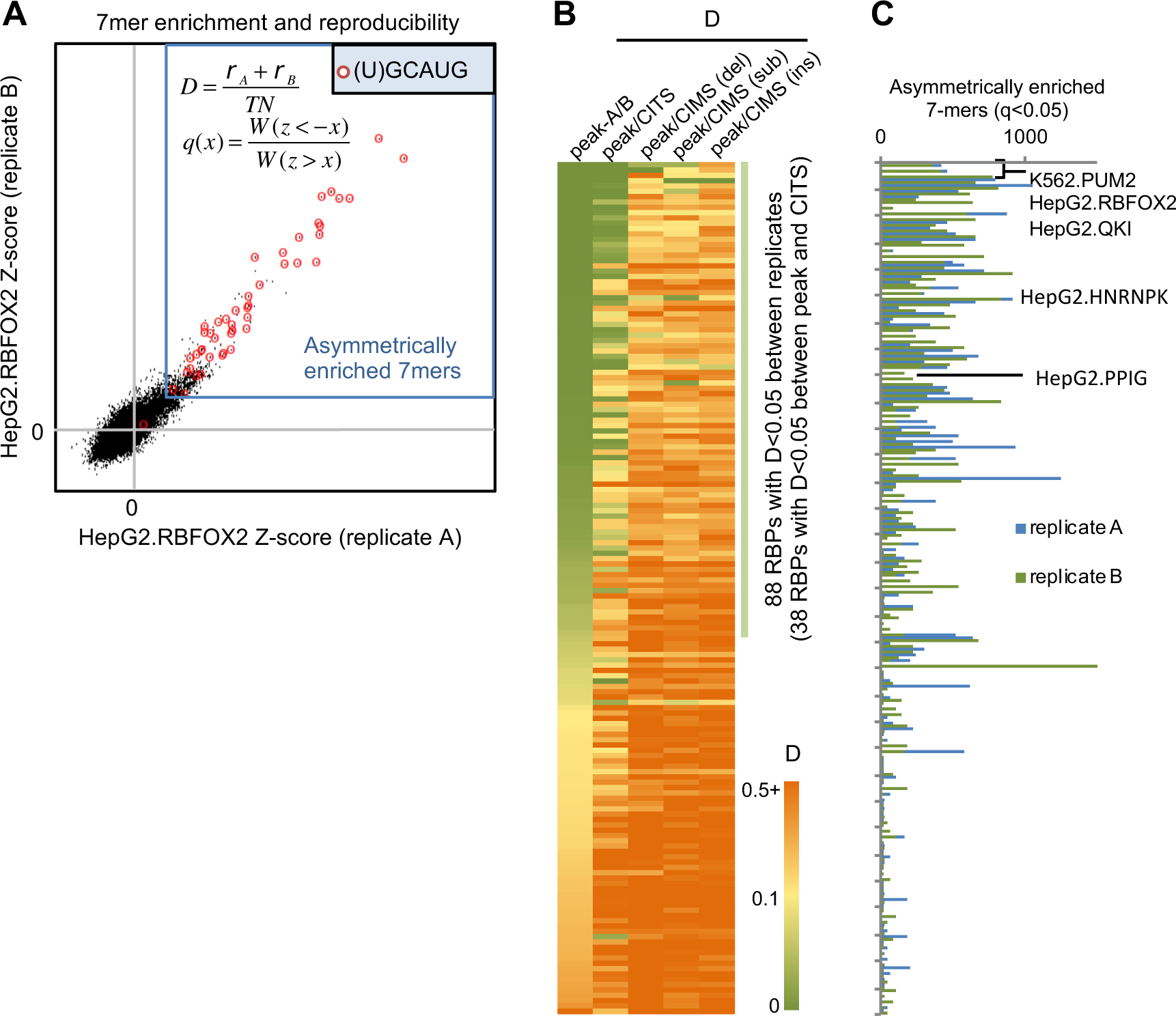
Quantitative measures used to characterize RBPs sequence specificity and reproducibility between replicate CLIP experiments. Related to Figures S3 and S4. (A) Illustration of top 7-mer disconcordance and asymmetric enrichment, using RBFOX2 eCLIP in HepG2 cells for an example. A z-score was calculated for each 7-mer based on its enrichment in CLIP tag cluster peaks for each replicate. 7-mers asymmetrically enriched in peaks are indicated using the blue box and 7-mers containing (U)GCAUG are highlighted in red. (B) RBPs are ranked based on the top 7-mer D-scores between peaks of the two replicates, between peaks and CITS, and between peaks and CIMS (deletions, substitutions and insertions analyzed separately). (C) Number of asymmetrically enriched top 7-mers for each RBP (q<0.05), shown in the same order as in (B).

Top 7-mers in 88/160 (55%) CLIP experiments showed D-score <0.05 (indicating that the top 20 7-mers in replicate A have an average ranking of 0.05 × 4^7^/2=~400 in replicate B and *vice versa*; Figure 2B). For RBPs with low D-scores between the two replicates in the same cell lines, they also have low D-scores when CLIP data from different cell lines were compared, indicating the same binding specificity in different cell types (Figure S4). In addition, RBPs with low D-scores in general have a larger number of significantly enriched 7-mers (Spearman ρ=−0.7, p<4.7e-25; Figure 2C). These observations together suggest that RBPs with low D-scores are more likely to have robust and reproducible binding specificity. While some RBPs with large D-scores between replicates could be due to technical issues, they might also tend to lack recognizable sequence specificity. In line with this notion, we found that a significantly higher portion of RBPs with D<0.05 have a RRM (36/88=41% vs. 27%; p=0.028; Fisher’s exact test), or K homology (KH) domain (14/88=16% vs. 1/72=1.4%; p=0.0018; Fisher’s exact test), compared to RBPs with D>0.05. This is presumably because these RNA-binding domains (RBDs) are well known to recognize specific sequences. Furthermore, a significantly higher portion of RBPs with D>0.05 do not have any annotated RBD (47/72=65% vs. 37/88=42%; p=0.0042; Fisher’s exact test).

We used the D-score metric to compare top 7-mers enriched in CLIP tag cluster peaks and those enriched in CITS or CIMS derived from different types of mutations to assess reliability of inferred crosslink sites. Among the 88 CLIP experiments with D<0.05 between replicates, 38 showed consistent 7-mer enrichment (D<0.05) compared to CITS, while few showed consistent 7-mer enrichment in CIMS. Based on these and other observations, we concluded that CIMS does not appear to provide reliable inference of crosslink sites in this dataset, and we focused on crosslink sites inferred by CITS for *de novo* motif discovery using mCross.

mCross was applied to CITS identified in the 160 CLIP experiments (with two replicates combined) and was able to discover one or more motifs for 144 experiments (the other 16 experiments do not show significantly enriched 7-mers, indicating lack of binding specificity). Overall, RBPs with low D-score between replicates tend to have more unambiguous motifs after similar ones were clustered together (Spearman rank correlation ρ=0.54, p=2.5e-12; Figure 3 and Table S2). An interactive, searchable web interface was also developed to facilitate access to this resource by the research community (http://zhanglab.c2b2.columbia.edu/index.php/MCross).

**Figure 3:**
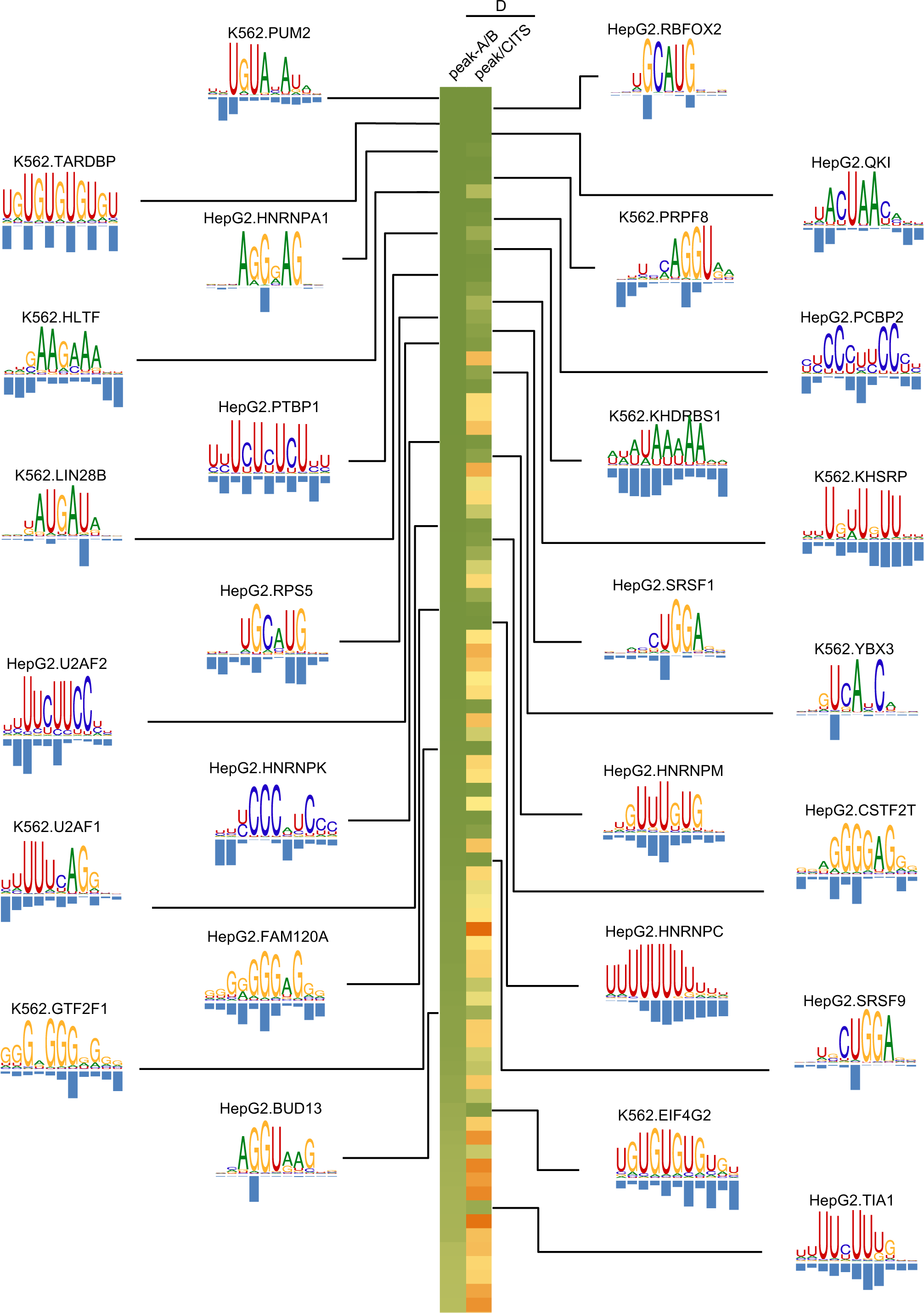
RBPs motifs and crosslink sites inferred by mCross. Results are shown for 38 experiments representing 27 distinct RBPs with D<0.05 between peaks of the two replicates and between peaks and CITS. Logos are only shown for the 27 distinct RBPs. For RBPs with multiple clusters of motifs, the representative motif with the highest likelihood score from the top cluster is shown.

### Validating RBP motifs by allelic protein-RNA interactions

A major challenge for *de novo* motif discovery is that an algorithm typically finds multiple motifs, leaving the user to decide which one is most reliable, if any. Mutagenesis, together with measurement of binding affinity or reporter assays, is the standard approach for experimental validation, but such validation is typically limited to a small set of selected binding sites, resulting in uncertainty in generalizability. We argued that protein-RNA interaction sites overlapping with SNPs represent a large number of natural perturbation experiments. In particular, the binding affinity of the two alleles at heterozygous SNPs can be directly compared by the allelic imbalance of CLIP tags (Figure 4A). We therefore performed allelic interaction (AI) analysis using heterozygous SNPs called from whole genome sequencing and eCLIP data (Figure S5 and Table S3; see STAR Methods for detail). In total, we identified 39,528 potential AI sites from HepG2 (an average of 565 sites per RBP) and 29,463 sites from K562 cells (an average of 331 sites per RBP; Table S4).

**Figure 4:**
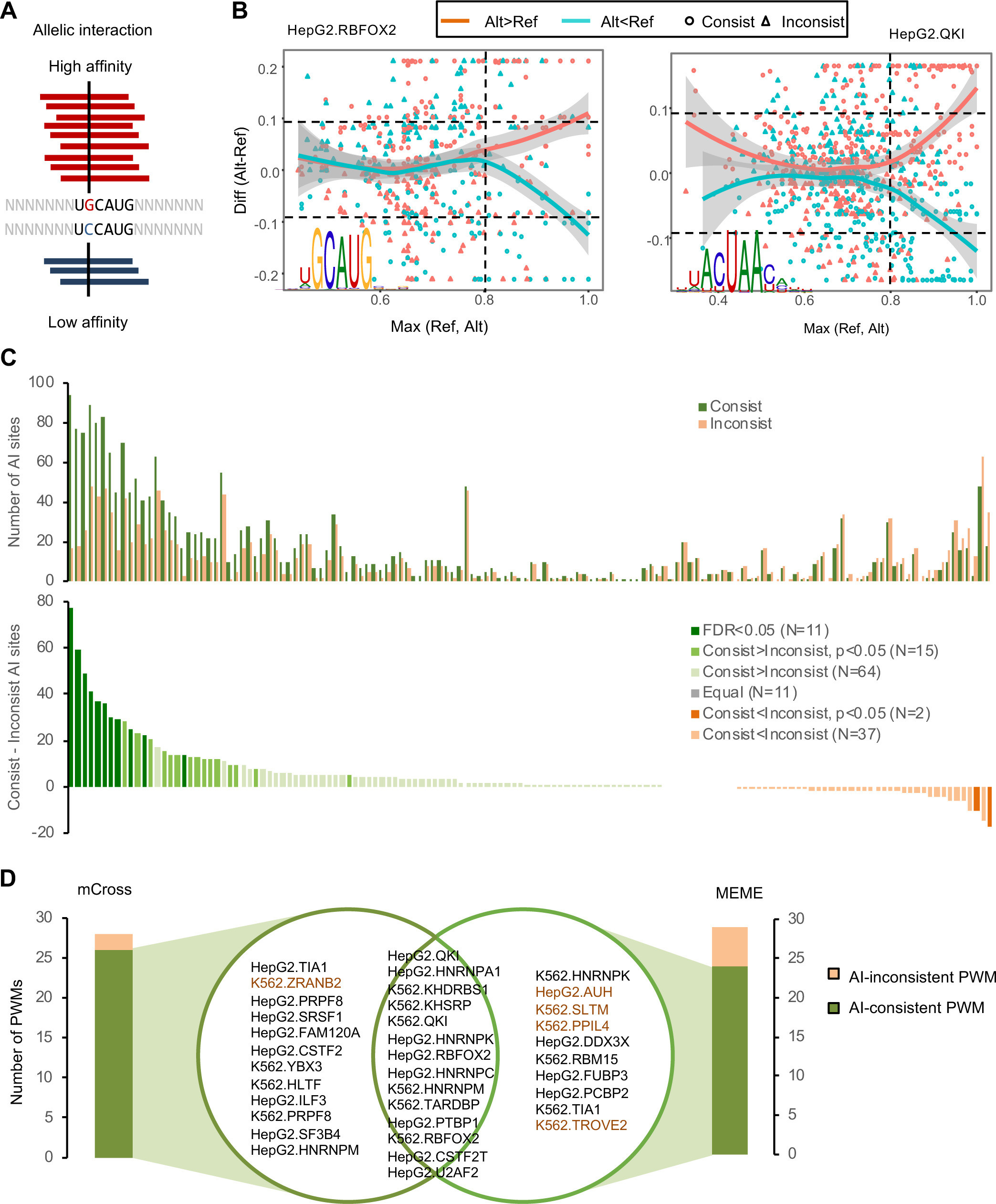
Evaluation of RBP motifs by allelic interactions. Related to Figure S5. (A) Schematic of a heterozygous SNP affecting Rbfox binding and the resulting allelic imbalance in eCLIP data. (B) SNPs showing allelic imbalance in RBFOX2 and QKI CLIP and the relationship between the allelic bias and motif score of the reference and alternative alleles. (C) The number of consistent and inconsistent AI sites for each RBP (top). A representative PWM is used for this analysis for each RBP. The excess of consistent over inconsistent AI sites is shown at the bottom and RBPs are color-coded based on the extent of excess using a Binomial test. (D) The list of AI-consistent PWMs identified by mCross and MEME using one representative PWM for each RBP by each method. The overlap of the two methods is shown. RBPs with D>0.05 between replicates are highlighted in red.

If the allelic imbalance detected in CLIP data is due to differential binding of the implicated RBP in the reference and alternative alleles, and the motif model accurately characterized the binding specificity of the RBP, one would expect that the allele with more CLIP tags would have a high motif score, and the other allele with fewer CLIP tags would have a reduced motif score. We denote these AI sites “consistent” sites (and otherwise inconsistent AI sites). For example, for the AI sites of RBFOX2 and QKI with more CLIP tags supporting the alternative allele (“red points”), the motif score is in general higher for the alternative allele, while for the AI sites with few CLIP tags supporting the alternative allele (“blue points”), the motif score is in general higher in the reference allele (Figure 4B). As expected, the trend is clear only for SNPs overlapping with a high-scoring motif site. For each RBP motif, we can thus obtain a subset of AI sites overlapping with high motif scores on either the reference or the alternative allele and also large motif score differences between the two alleles. The proportion of consistent AI sites was estimated as a measure of the accuracy of the motif model.

We initially used a representative PWM (the first PWM discovered by mCross) for each RBP to compare with AI sites to avoid any bias, as selection of the PWM is independent of AI site analysis. For a majority of RBPs, consistent AI sites are in excess as compared to inconsistent AI sites, while a much smaller number of RBPs show the opposite pattern (90 vs. 39 RBPs; Figure 4C). Using a more stringent threshold (p<0.05; Binomial test), 26 RBPs have significantly more consistent AI sites compared with inconsistent AI sites (denoted AI-consistent PWMs) and only 2 RBPs show the opposite pattern (Figure 4C). For the remaining RBPs, which individually have an insufficient number of AI sites for statistical analysis, the overall proportion of consistent AI sites is also significantly higher than 0.5 when they were analyzed in aggregate (p<0.004, Binomial test). Therefore, the concordance between the allelic imbalance of CLIP tags and changes in motif scores provides an unbiased validation that the motifs defined by mCross reliably reflect RBP binding specificity. Furthermore, these AI sites also provide a list of genetic variations in the human populations that directly affect protein-RNA interactions with potential impact on downstream post-transcriptional gene expression regulation and phenotypes.

### Comparison of mCross with other methods by allelic interaction sites

We next used AI site analysis to evaluate PWMs derived by other methods to provide an unbiased comparison of these methods and mCross. First, we compared mCross with MEME, a widely used program for *de novo* motif discovery (Bailey and Elkan, 1994). As MEME discovered multiple motifs for each RBP, we used the top motif for each RBP for comparison. Among the 159 PWMs derived from HepG2 and K562 data by MEME, 24 are AI-consistent PWMs and 5 are AI-inconsistent PWMs (p<0.05; Binomial test), as compared to 26 AI-consistent PWMs and 2 AI-inconsistent PWMs derived by mCross using the same criteria (Figure 4D). If AI-consistent PWMs are more reliable, we expect them to have low top 7-mer D-scores between CLIP replicates. Indeed, for the 14 RBPs with AI-consistent PWMs identified by both mCross and MEME, all have D<0.05. Similarly, 11/12 RBPs with AI-consistent PWMs only identified by mCross have D<0.05. On the other hand, 4/10 RBPs with AI-consistent PWMs only identified by MEME have D>0.05. Therefore, motifs derived by mCross showed better concordance with AI sites than MEME, suggesting they characterize RBP binding specificity more accurately.

We also compared 18 RBPs with both eCLIP and RNAcompete data (Ray et al., 2013). Among them, 6 RBPs have AI-consistent PWMs derived from RNAcompete, and 10 RBPs have AI-consistent PWMs derived by mCross. For the four RBPs with AI-consistent PWMs only by mCross (HNRNPA1, TARDBP, TIA1, and U2AF2), all have well characterized binding specificity.

Given the effectiveness of AI sites for comparison of different PWMs of the same RBPs, we selected the optimal motifs for each RBP based on their consistency with AI sites (Table S5). This analysis allowed us to obtain a final list of 16 RBPs in HepG2 and 13 RBPs in K562 with AI-consistent PWMs (FDR<0.1; Figure 5) and a subset of AI sites filtered by these PWMs that most likely affect protein-RNA interactions directly (Table S4).

**Figure 5:**
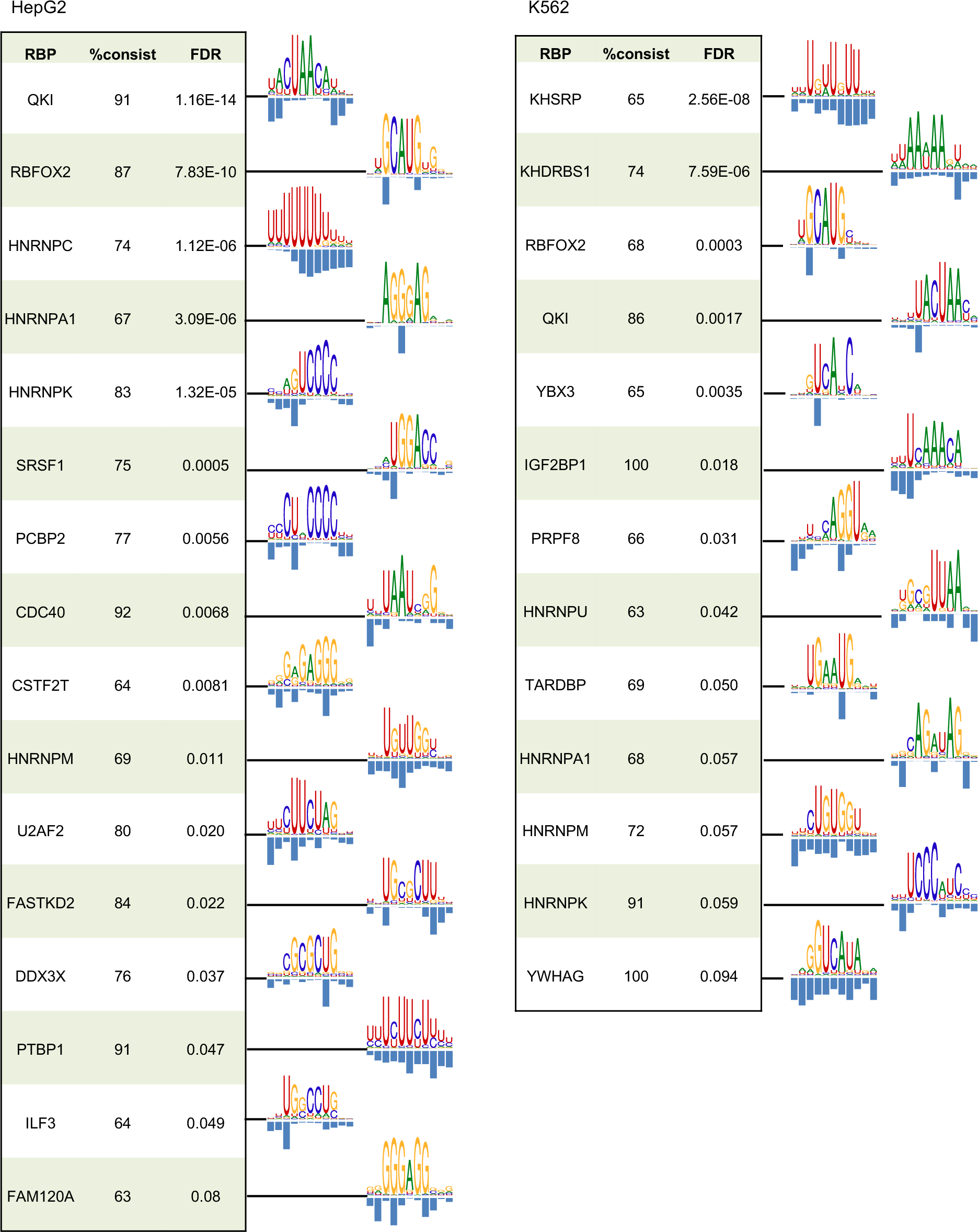
The list of optimal AI-consistent PWMs discovered by mCross. Only RBPs with AI-consistent PWMs at FDR<0.1 are shown.

### SRSF1 recognizes GGA clusters to activate exon inclusion

Overall, for well characterized RBPs, the motifs defined by mCross agree very well with the previously defined motifs (Figures 3 and 5). There are also interesting exceptions. For example, we found a (U)GAU motif for LIN28, which is distinct from the previously characterized GGAG motif. mCross also determined that SRSF1 binds a (U)GGA motif, which is the half site of the canonical GGAGGA motif. The functional significance of the LIN28 motif was described in our recent study (Ustianenko et al., 2018). Here we focus on the importance of the GGA motif for SRSF1-dependent alternative splicing.

SRSF1 is the founding member of the SR protein family, which is important for regulation of both constitutive and alternative splicing. mCross identified a UGGA motif for SRSF1 with predominant crosslinking in the U1 position. The GGA motif represents a half site of the previously defined SRSF1-binding consensus GGAGGA from SELEX and CLIP data (Sanford et al., 2009; Tacke and Manley, 1995) (Figure 6A). The first nucleotide of the UGGA motif likely reflects crosslinking bias, as SNPs at this position do not affect binding, while the other three positions are important (Figure S6A). Interestingly, a previous structural study suggested that the second RRM of SRSF1 directly contacts a GGA half site (Figure 6B) (Clery et al., 2013), which agrees well with the motif discovered by mCross.

**Figure 6:**
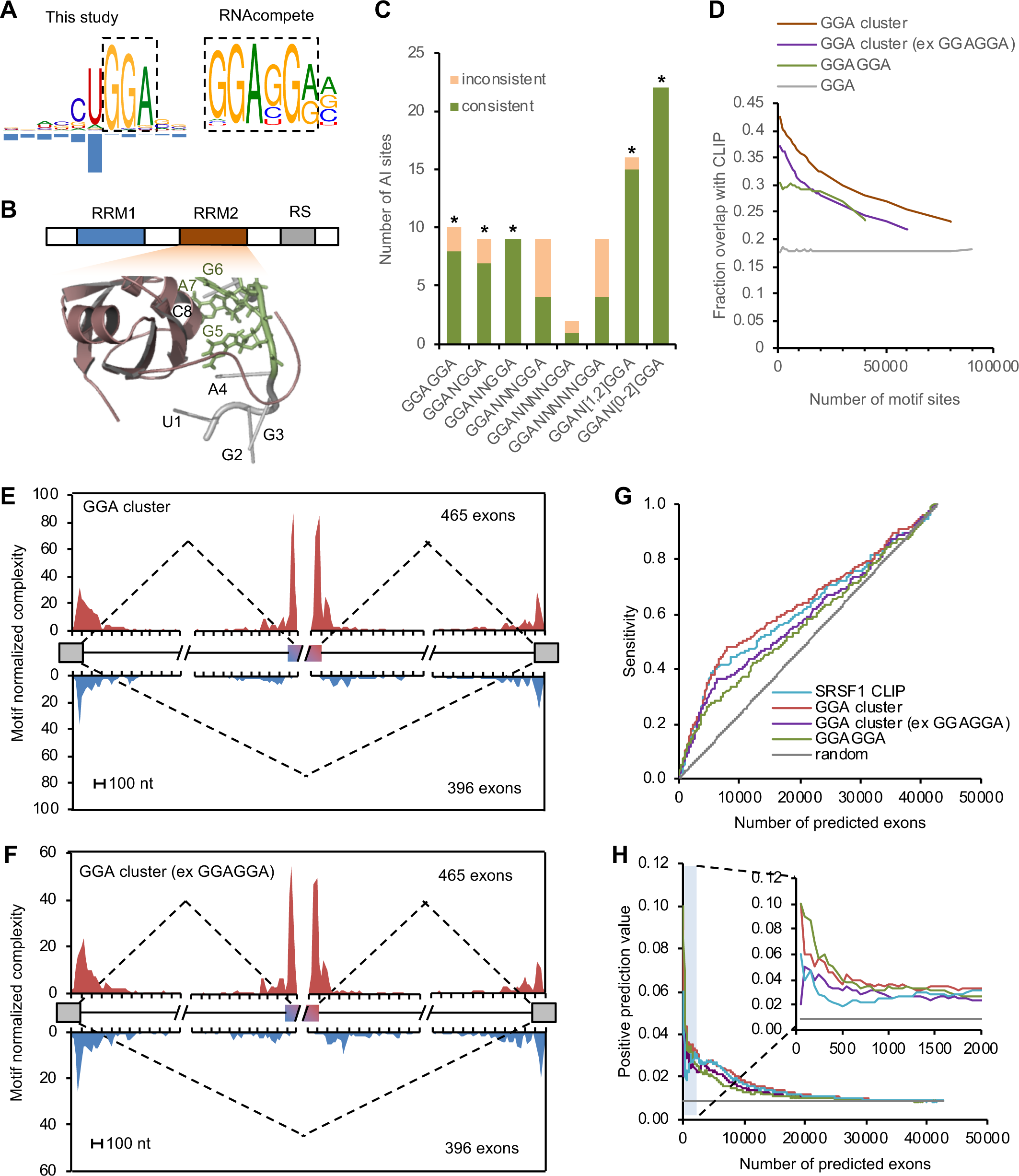
SRSF1 recognizes a GGA motif in vivo sufficient for regulation of alternative splicing. Related to Figures S6 and S7. (A) SRSF1 binding motif identified by mCross (GGA) and by RNAcompete (GGAGGA). (B) NMR structure of SRSF1 RRM2-RNA complex (PDB accession: 2m8d). The GGA half site directly contacting the RRM is highlighted in green. (C) Allelic interaction analysis using bipartite GGA-N[_02_]-GGA motif. Binomial test was used to test whether the excess of consistent AI sites over inconsistent AI sites is significant (* p<0.05). (D) Overlap of predicted GGA clusters with SRSF1 eCLIP tag clusters at different ranks of motif scores. GGA clusters without overlapping with GGAGGA and conserved GGAGGA sites ranked by BLS were used for comparison. Only motif sites in the CDS region were used for this analysis. (E, F) RNA map showing GGA clusters enriched in cassette exons with SRSF1-dependent inclusion, but depleted in alternative exons with SRSF1-dependent skipping. Results were obtained for all GGA clusters (E) or GGA clusters without overlapping with GGAGGA (F). (G, H) Prediction of SRSF1-activated cassette exons using GGA clusters. Conserved GGAGGA sites and CLIP tag clusters are used for comparison. The sensitivity (G) or positive prediction value (H) with respect to varying number of predicted exons by each method are shown.

Since the GGA motif alone has very limited information content, we reasoned that sufficient targeting specificity for SRSF1 has to be achieved by binding to a cluster of GGA elements near each other. Mechanistically, GGA clusters can be bound by multimerization of multiple SRSF1 proteins with each RRM contacting one GGA motif site (Liu et al., 1998). To test this hypothesis, we first searched a bipartite GGA-N_x_-GGA motif with a spacer. Indeed, we found AI sites overlapping GGAN_[1–2]_GGA also showed an excess of consistent sites, similar to the pattern found for the canonical motif GGAGGA (Figure 6C). We therefore predicted GGA clusters using mCarts, which integrates the number of GGA elements, their spacing, conservation, and accessibility as determined by predicted RNA-secondary structures (Weyn-Vanhentenryck and Zhang, 2016; Zhang et al., 2013) (Figure S6B). GGA clusters predicted using the models trained by HepG2 and K562 CLIP data are highly similar to each other qualitatively and quantitatively (Figure S6C, D). Therefore, GGA clusters trained on HepG2 CLIP data were used for detailed analysis described in this study.

The predicted GGA clusters with higher motif scores have a higher overlap with CLIP tag clusters (up to over 40%, as compared to 18% overlap observed from individual GGA elements; Figure 6D). This is true after excluding clusters containing GGAGGA, suggesting SRSF1 binds to GGA clusters without requiring GGAGGA on a genome-wide scale.

To test whether the predicted GGA clusters are sufficient to regulate alternative splicing, we identified cassette exons showing altered splicing upon SRSF1 knockdown, using RNA-seq data generated by ENCODE (Van Nostrand et al., 2017). As a positive control, we first generated an RNA map that predicts the impact of SRSF1 binding position on splicing using CLIP tags and predicted GGAGGA motif sites. As expected, substantial enrichment of SRSF1 binding was observed in alternative exons with SRSF1-dependent inclusion, and depletion of SRSF1 binding was observed in cassette exons with SRSF1-dependent skipping (Figure S7A, B), which is consistent with the known role of SRSF1 in activating exon inclusion by binding to exon splicing enhancers (ESEs). Importantly, the same pattern was obtained using predicted GGA clusters, even after excluding GGA clusters overlapping with GGAGGA (Figure 6E, F). These results suggest that the predicted GGA clusters without GGAGGA are functional in activating exon inclusion.

We next evaluated how accurate GGA cluster motif score predicts individual SRSF1 target exons. To this end, we scored every exon by using the strongest GGA cluster in the alternative exon. Exons ranked by conserved GGAGGA using branch length score (BLS) and CLIP tag cluster scores were used for comparison. Among exons with SRSF1-dependent inclusion in both HepG2 and K562 cells (ΔΨ>0.1, and FDR<0.05), 54% (200/373) have predicted GGA clusters, as compared to 21% among all cassette exons. The majority of SRSF1-dependent exons harboring GGA clusters (137/200=69%) do not have GGAGGA. We ranked the exons by their score and calculated the sensitivity and positive prediction value (PPV) of predicting SRSF1-activated exons captured at each rank. This allowed us to compare the performance of GGA clusters and GGAGGA in determining SRSF1-dependent splicing regulation. We found that the GGA clusters are more predictive than conserved GGAGGA motif sites, as reflected in increase in both sensitivity and PPV (Figure 6G, H). Importantly, the performance of the GGA clusters is similar to, if not higher than, that of CLIP cluster scores and has more scored exons, indicating that the GGA clusters are both reliable and able to complement the CLIP data. Excluding GGA clusters overlapping with GGAGGA remains predictive of SRSF1-dependent exons, despite a minor reduction in performance.

We also reasoned that SRSF-dependent exons identified through knockdown experiments might represent an underestimation of the contribution of SRSF1 in splicing regulation due to compensation by other SR proteins or other mechanisms. To address this issue, we examined whether GGA clusters are predictive of exon inclusion level. Indeed, exons with high inclusion are much more enriched in predicted GGA clusters, even after exclusion of GGAGGA. In HepG2 cells, 30.2% cassette exons with inclusion level Ψ≥0.9 have predicted GGA clusters, as compared to 8.8% for cassette exons with Ψ ≤0.1, suggesting a conservative estimate that over 20% cassette exons with high inclusion are regulated by SRSF1. About 80% of these GGA clusters do not overlap with GGAGGA and similar results were obtained when more stringent thresholds on the motif score were used (Figure S7C, D). Altogether, our analysis suggests that SRSF1 has a much larger repertoire of transcripts that it can recognize to regulate their splicing.

As particular examples, we found that SRSF1 regulates a cassette exon in both *HNRPD* (ΔΨ=0.34, FDR=1.9e-198) and *HNRPDL* (ΔΨ=0.64, FDR=1.9e-159; Figure 7A, B) (these changes are in K562 cells; consistent changes in HepG2 cells, although somewhat smaller in magnitude). In both cases, GGA clusters were predicted in the alternative exon, supported by robust CLIP tag clusters. There are two GGA clusters in HNRNPDL, without any GGAGGA sites, suggesting the importance of GGA half sites for SRSF1 binding and splicing regulation. The GGA cluster in HNRPD has a GGAGGA motif with four additional GGA sites separated by variable number of nucleotides. Interestingly, in each case the alternative exon encodes an intrinsically disordered peptide enriched in a glycine and tyrosine (GY) dipeptide motif that mediates multivalent hnRNP assemblies with global impact on downstream splicing regulation (Gueroussov et al., 2017). Our results suggest that SRSF1 serves as an important upstream modulator of this mechanism.

**Figure 7:**
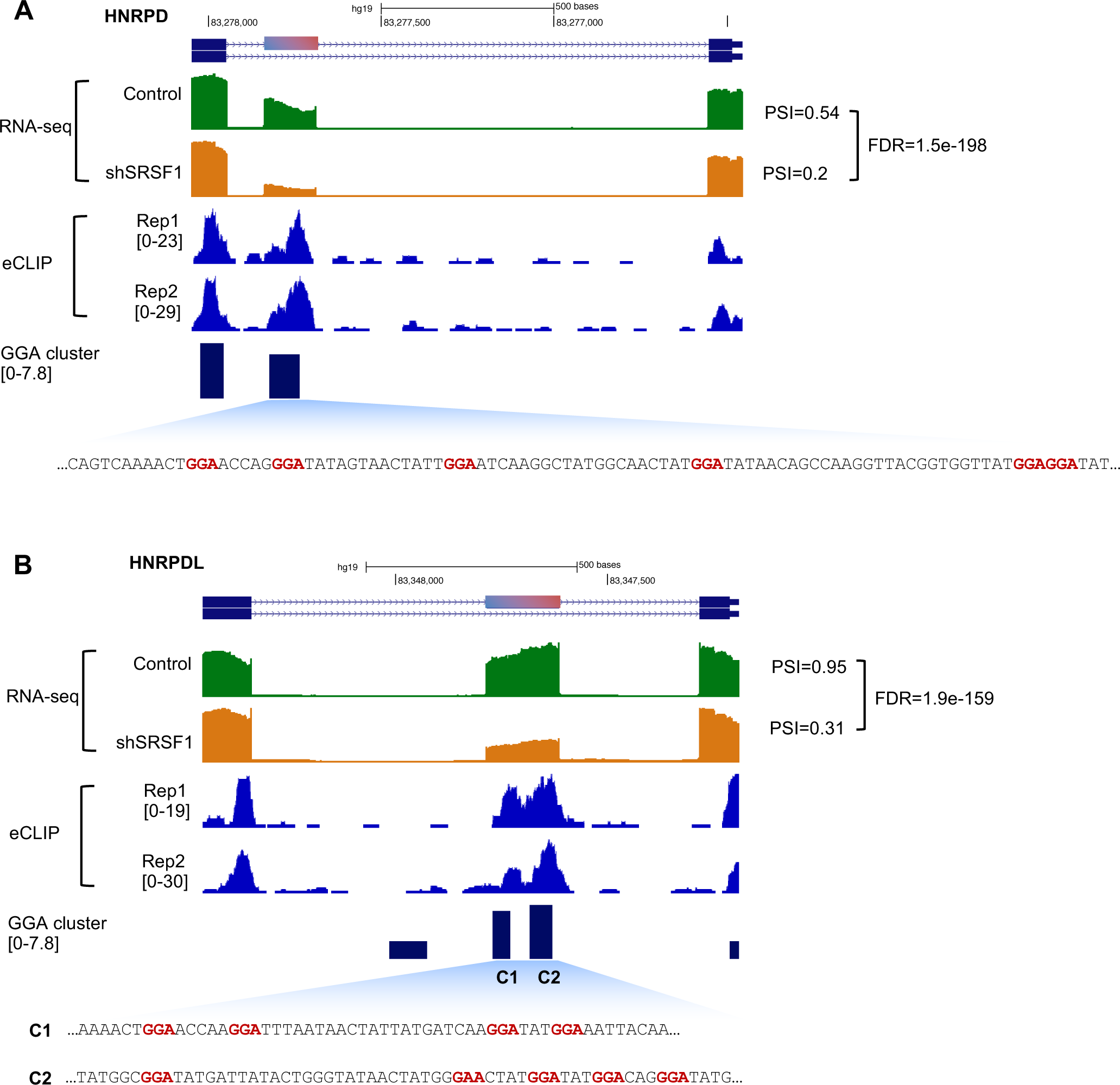
SRSF1 regulates a cassette exon in HNRPD and HNRPDL through GGA clusters. (A) HNRPD. (B) HNRPDL. In each case, the cassette exon encodes an intrinsically disordered peptide enriched in glycine-tyrosine (GY) motif mediating multivalent hnRNP assemblies. RNA-seq data with and without SRSF1 knockdown and SRSF1 eCLIP data in K562 cells are shown.

## Discussion

The intrinsic flexibility of RBPs in recognizing their RNA regulatory sequences imposes a big challenge in accurate characterization and predictive modeling of their specificity, even when a large number of binding footprints are mapped by CLIP. To address this problem, we developed a new statistical model for *de novo* motif discovery using CLIP data named mCross. mCross builds on the critical observation that protein-RNA crosslinking in CLIP experiments frequently occurs at specific positions within the motif, which can be mapped at single nucleotide resolution. These crosslink sites provide precise registers of motif sites in input sequences, and thus dramatically reduce the search space during *de novo* motif discovery. We note that the proximity of RBP motif sites to crosslinking events was previously used in another algorithm Zagros to facilitate motif discovery using a different statistical model (Bahrami-Samani et al., 2015). The Zagros model does not distinguish crosslink propensity at different positions in the motif. In addition, crosslinking events were defined in individual CLIP tags, which are noisy due to technical issues. When PAR-CLIP data were used, crosslink sites at 4SU might not reflect direct-protein contact in the native conditions. Therefore, to our knowledge, mCross is the first model to jointly model RBP sequence specificity and the precise protein-RNA crosslink sites at specific motif positions at single-nucleotide resolution.

We applied mCross to the largest CLIP datasets generated thus far by ENCODE to define motifs of 112 unique RBPs. Importantly, we developed multiple quantitative measures to assess the reliability of the results. We performed genome-wide AI site analysis using CLIP to detect SNPs affect protein-RNA interactions, as these AI sites provide a large number of naturally occurring perturbation experiments *in vivo* that can be used to validate the accuracy of discovered motifs. Our analysis suggests mCross performs favorably compared to other state-of-the-art methods. In addition, when multiple, distinct motifs were discovered for the same RBP, in which ambiguity frequently arises, AI analysis also provides a means of selecting the most reliable motif. On the other hand, AI sites filtered by PWMs also provide a subset of high-confidence SNPs that directly affect protein-RNA interactions, with potential functional implications in human populations.

We expect the resulting motifs, complemented by an interactive, searchable web interface, will be a useful resource for the research community to make new discoveries. In a recent study, we showed the importance of a novel LIN28 (U)GAU motif discovered by mCross in differential binding by LIN28 and suppression of two subclasses of let-7 microRNAs that are major downstream targets of LIN28 (ref.(Ustianenko et al., 2018)). In this case, crosslinking of LIN28 to the last uridine in the motif was also experimentally validated(Ransey et al., 2017). In this study, we found SRSF1 binds a cluster of GGA half sites as the predominant mode of binding besides the previously characterized GGAGGA element, and this flexibility leads to a much larger repertoire of target transcripts that were not previously appreciated. These case studies exemplify how improved definition of RBP binding specificity can lead to mechanistic insights into RNA regulation.

## Acknowledgements

We thank members of the Zhang laboratory for helpful discussion. This study was supported by grants from the National Institutes of Health (NIH) (R01GM124486 and R03HG009528 to CZ). SB was in part supported by a Columbia Precision Medicine Research fellowship. High-performance computation was supported by NIH grants S10OD012351 and S10OD021764.

## Author Contributions

HF, SB, and CZ conceived the study; HF, SB, SMW, JW, AS, EDF and CZ performed data analysis; AK designed the web interface. HF, SB and CZ wrote the paper with input from all authors.

## Materials and Methods

### CLIP data processing

We obtained CLIP data for Rbfox1–3 (Weyn-Vanhentenryck et al., 2014), Nova (Zhang et al., 2010), Ptbp2 (Licatalosi et al., 2012), Mbnl2 (Charizanis et al., 2012), and Lin28A (Cho et al., 2012) from separate studies. We previously analyzed Rbfox, Ptbp2 and Mbnl2 CLIP data part of the original studies using the CIMS package (Moore et al., 2014), a predecessor of CLIP Tool Kit (CTK) (Shah et al., 2017). Lin28A CLIP data was analyzed by the same pipeline (Moore et al., 2014). For each dataset, unique CLIP tags and mutations in unique tags, originally derived based on mapping to mm9, were liftOver to mm10. For CIMS analysis of deletions, we only included single-nucleotide deletions (and excluded deletions of two or more consecutive nucleotides).

eCLIP data of 70 RBPs in HepG2, 89 RBPs in K562, and 1 RBP in adrenal gland were downloaded from the ENCODE website (https://www.encodeproject.org; as of Dec 30, 2016) (Van Nostrand et al., 2017; Van Nostrand et al., 2016). In total, this dataset is composed of 112 unique RBPs, with 47 RBPs assayed in both HepG2 and K562 cells (Table S1). All mock control (input) data were also downloaded. The raw reads were processed to obtain unique CLIP tags mapped to hg19 using CTK (Shah et al., 2017), as described previously (Ustianenko et al., 2018). Only read2 (the read starting from 5’ end of the RNA tag) was used for analysis described in this paper. For each RBP, unique tags from the two replicates were combined for all analyses, except for evaluating reproducibility between the two replicates (see below). Significant CLIP tag clusters were called by requiring P<0.001 after Bonferroni multiple-test correction. Crosslinking-induced truncation sites (CITS) were called by requiring FDR<0.001. Crosslinking induced mutation sites (CIMS) were also examined but not reported in this paper because they appear to have low signal-to-noise ratio.

### 7-mer enrichment analysis

To provide seeds for *de novo* motif discovery using mCross, we performed 7-mer enrichment analysis using significant peaks with peak height (PH)≥10 tags. Peaks were extended for 50 nt on either side relative to the center of the peak to extract the foreground sequences. Background sequences were extracted from the flanking regions of the same size (−550, −450) and (450, 550) relative to the peak center. Sequences with more than 20% of nucleotides overlapping with repeat masked regions were discarded. 7-mers were counted in repeat-masked foreground and background sequences, and the enrichment of each 7-mer in the foreground relative to the background was evaluated using a binomial test. A z-score (and a p-value) was derived for each 7-mer, and denoted raw z-score.

The raw z-scores were then normalized because we noticed a general enrichment of certain 7-mers (such as G-rich elements) in many experiments. To minimize potential experimental biases and non-specific protein-RNA interactions, we normalized the raw z-score of each 7-mer by subtracting median across all experiments followed by scaling using the median absolute deviation (MAD), a robust estimate of the standard deviation. The resulting score was denoted the normalized z-score, and was used to rank and identify top 7-mers.

We developed an asymmetric enrichment score for each top 7-mer to evaluate their statistical significance, as an assessment whether an RBP has binding specificity. We argue that if there is no 7-mer that is significantly more enriched in the foreground than the background, the distribution of the normalized z-scores should be symmetric. On the other hand, if an RBP shows high affinity to (a relatively small subset of) specific 7-mers, it should show a heavy tail on the right side of the distribution. We therefore derived a false discovery rate (FDR) based on the symmetry of the null distribution.

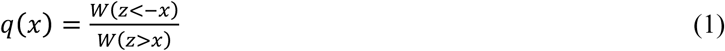

for each z=x(>0). W(.) denotes the number of 7-mers satisfying the specified criterion.

### The mCross model and the optimization algorithm

Most of the current *de novo* motif discovery tools (such as MEME (Bailey and Elkan, 1994) and HOMER (Heinz et al., 2010)) use a standard model of a position-specific weight matrix (PWM) to characterize the specificity of DNA- or RNA-binding proteins. Given a set of sequences bound by a specific RBP containing *K* binding sites of width *W*, the likelihood ratio of observing the data based on the motif model versus the background model can be written as follows:

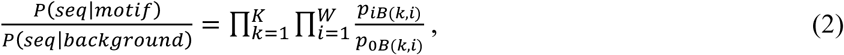

where *p_iB_*_(_*_k, i_*_)_ and *P*_0_*_B_*_(_*_k, i_*_)_ are the probability of observing base *b*= *B*(*k, i*) in position *i* of site *k* according to the motif and background models, respectively.

After log transformation and simple rearrangements,

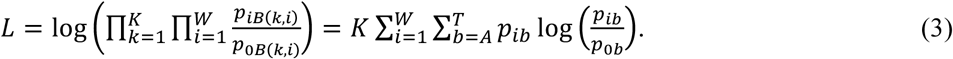

The mCross model augments the standard PWM model by jointly modeling the RBP sequence specificity and the precise protein-RNA crosslink sites at specific motif positions at single-nucleotide resolution. Denote *q_i, k_* and *q*_0_ the probability of protein-RNA crosslinking at position *i* of the motif site *k* according to the motif and background models, respectively, the likelihood ratio of observing *K* sites of size *W* can be written as:

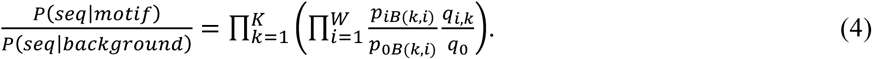

After rearrangement, the log likelihood ratio can be written as follows:

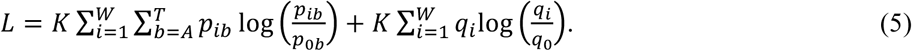

This model can be extended to allow crosslinking in specific positions outside the core motif:

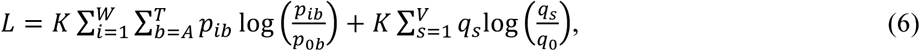

where *s* = 1,2,…, *V* indicates positions in the core motif or immediate flanking sequences (e.g., 2-nt extension on both sides of the core motif).

To search for parameters *p_ib_* and *q_i_* that optimize the objective function (we assume the background probabilities *p_0b_* = 0.25 and *q*_0_ = 1/*V* in this study), mCross currently uses a seed-based search strategy(Sahin and Sur, 2015). In brief, a ranked list of top 7-mers is obtained based on their normalized z-score and asymmetric enrichment score (q<0.05; we limit to the top 10 if there are more than 10 7-mers with q<0.05). RBPs without asymmetrically enriched 7-mers were not analyzed.

To initiate motif search, we first grouped top 7-mers with ≤2 mismatches without allowing shifts; each 7-mer group initiates one motif. Specifically, each 7-mer in a group is extended with the degenerate nucleotide ‘N’ for 2 nt on each side and then used as a seed to search for exact or inexact matches (≤*m* mismatches; *m*=1 for this study) around crosslink sites derived from CIMS or CITS analyses. These matches provide the list of all candidate RBP binding sites. Initially, all candidate sites are included to derive the motif model and calculate the log likelihood ratio *L*. An iterative procedure is then used to exclude or include each candidate site based on whether the adjustment improves the likelihood. The algorithm stops upon convergence or reaching the maximum iterations.

The objective function in eq. (6) in general favors degenerate motifs when more than one site is allowed in each input sequence. To reward more specific motifs, we introduced modifications to eq. (6) to generate results presented in this paper:

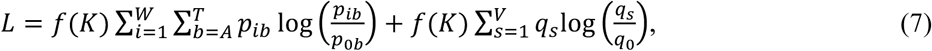

where 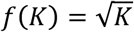.

Among the 160 eCLIP experiments (with replicates combined), mCross discovered at least one motif for 144 experiments.

### Motif clustering

For each RBP, we clustered similar motifs reported by mCross using Stamp (Mahony and Benos, 2007). PWMs were trimmed from both sides to remove positions with low information content ≤0.2, where information content was defined as in (Schneider et al., 1986). Pearson correlation coefficient was adopted to measure the distance of each compared pair of PWMs. Local Smith-Waterman ungapped alignment and unweighted pair group method with arithmetic mean (UPGMA) were used to perform alignment and grow the cluster tree. The number of clusters was determined by minimizing the Calinski-Harabasz (CH) index provided by Stamp. If there is no global optimized cutoff using CH index, the clustering trees were cut at height 0.05.

### Reproducibility of RBP sequence specificity

We developed rank-based measure of disconcordance of top 7-mer enrichment between the two replicates. To this end, we derived normalized z-score to rank 7-mers for each individual replicate. For the top *T* of a total of *N*=4^7^=16,384 7-mer (*T*=20 for this study) with ranksum *c*= *T*(*T* + 1)/2 in replicate A, we obtained their rank sum in replicate B:

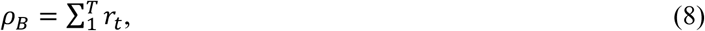

where *r_t_*(*t* = 1,…, *T*) is the rank of each top 7-mer.

Vice versa, for the top *T* 7-mer in replicate B, we obtained their rank sum *ρ_A_* in replicate A. The disconcordance of the two replicates is measured by a score *D*.

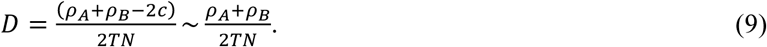

We can similarly compare whether top 7-mers identified at CLIP tag cluster peaks are also ranked high in sequences near crosslink sites. Denote the rank sum of the top 7-mers in sequences of crosslink sites *ρ*.

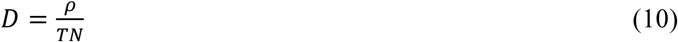

In this work, we consider an RBP with *D*<0.05 between the two replicates as having reproducible sequence specificity. We also used D-score to compare top 7-mers in peaks and CITS, and to compare CLIP and RNAcompete data (Ray et al., 2013).

### SNP calling in HepG2 and K562 cells using eCLIP and whole genome sequencing data

If a protein-RNA interaction site is affected by genetic variation at a heterozygous SNP site, i.e., allelic interaction (AI), the two alleles will have different numbers of supporting CLIP tags. We used global analysis of AI sites to validate RBP motifs discovered by mCross.

To identify AI sites in CLIP data, we first called heterozygous SNPs in HepG2 and K562 cells using eCLIP (including mock) and whole genome sequencing (WGS) data. For each sample (either CLIP or mock), the genomic mapping information of the unique tags was extracted, stored in a sam file, and converted to bam using SAMtools (Li et al., 2009). The bam files of all samples (including both CLIP and mock data) of the same cell line were merged together for HepG2 and K562, respectively. The variant-calling procedure was implemented following GATK (v3.8)’s best practice recommendations for RNA-seq data with minor modifications (McKenna et al., 2010). In particular, no “MarkDuplicate” step was carried out as PCR duplicates have already been removed using a method optimized for CLIP data and the resulting unique tags were used as input. Per-base sequencing error was estimated by “BaseRecalibrator”. Then, we separately called variants for each cell line with “HaplotypeCaller” (with stand_call_conf set to 20 to ensure high sensitivity). Limited by the lack of truth/training sets required by the Variant Quality Score Recalibration (VQSR) step for eCLIP data, we adopted hard filters for variant filtration. Only bi-allelic SNPs with QualByDepth (QD) > 5 and depth (DP)>10 were kept. SNPs overlapping with RNA editing sites, as annotated in DARNED(Kiran and Baranov, 2010), were excluded.

To complement and improve genotype calls derived from eCLIP data, we performed variant calling using WGS data of HepG2 and K562 cells (Table S3), respectively, following the GATK “best practices” protocol for DNA-seq data. Briefly, for each cell line, WGS reads from different platforms were aligned to hg19 using BWA (Li and Durbin, 2010) and were merged together. We performed “MarkDuplicates” to remove PCR duplicates and used “BaseRecalibrator” to ensure the base quality. The variant calling step was carried out by “HaplotypeCaller” (with stand_call_conf set to 30). For variant filtering, we excluded variants on annotated RNA editing sites in DARNED (Kiran and Baranov, 2010) and applied the VQSR step to ensure that 99% of Hapmap SNPs in our data were included in the final call set. Only WGS SNPs covered by ≥10 eCLIP reads were used for genotype correction. We defined three subsets of SNPs to be considered in the following AI site analysis: 1) SNPs called heterozygous consistently in both eCLIP and WGS data. 2) the intersection of eCLIP and WGS data, but as heterozygous only in WGS data (the genotype call from WGS data was used for these SNPs); and 3) heterozygous SNPs called only in eCLIP data. We also filtered the last two categories by keeping only bi-allelic SNPs. SNPs called only from eCLIP data that were called as homozygous in WGS data or were inconstant with dbSNP (v138) genotypes were also excluded.

Finally, for each cell line and RBP, the number of unique CLIP tags supporting each allele of a heterozygous SNP was extracted from bam files with SAMtools (Li et al., 2009). We inferred the sense transcript strand of each SNP by pooling CLIP tags from all CLIP and mock data and counting the number of tags from each strand. The strand with a majority of supporting tags was considered the sense strand. Only heterozygous SNPs with unambiguously inferred transcript strand (i.e., #sense read/(#sense read+#antisense read)>0.9) were included in our analysis. The final dataset consisted of 229,265 and 155,388 heterozygous SNPs in HepG2 and K562 cells, respectively (Table S3).

### Identification of allelic interaction sites from eCLIP data

To assess allelic binding of each RBP at a heterozygous SNP, we counted the number of sense CLIP tags of the RBP supporting each allele. Two types of control data were used for comparison: 1) pooled CLIP data of all other RBPs (except the RBP under consideration) in the same cell line (e.g., RBFOX2 vs. CLIP data of all other RBPs except RBFOX2); 2) all pooled mock data in the same cell line (e.g., RBFOX2 vs. mock). For each comparison, the magnitude of allelic imbalance was defined as |Δ*A*|=|*AAF*_RBP_-*AAF*_control_|, where AAF (alternative allele frequency) was estimated from the number of sense CLIP tags supporting the alternative allele divided by the total number of sense CLIP tags overlapping with the SNP. The statistical significance of AI was evaluated using a Fisher’s exact test. For this analysis, we considered all heterozygous SNPs with coverage ≥10 sense tags. Sites with 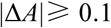 and p<0.05 in either of the two comparisons were called significant AI sites (Table S4).

### Validation of PWMs using AI sites

For each PWM derived by mCross, we trimmed the flanking motif positions with information content≤0.4. For each AI site, the PWM score (Liu and Stormo, 2005) of the sequence associated with allele was calculated. Briefly, the reference and alternative allele sequences flanking each AI site of size 2*W-*1 were extracted, where *W* is the width of the trimmed PWM. We scanned the sequences and calculated the PWM scores of all possible motif sites:

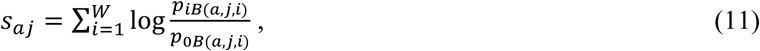

where *a* indicates the reference or alternative allele, *j* = 1, 2, *…, W* is the offset of the motif site and *i* is the position of the nucleotide in the motif site. For the background base composition, we used *p*_0*B*(*a, j, i*)_ = 0.25.

### The PWM score was then normalized

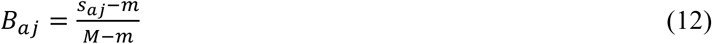

where *M* and *m* are the maximal and minimal possible scores of the PWM. The binding affinity of the RBP to allele *a* is estimated to be *B_a_ =* max*_j_ B_aj_*. The SNP is considered to affect RBP binding if max(*B_Ref_, B_Alt_*) > *α* and |*B_Ref_ − B_Alt_| > δ*, where *α* and *δ* are thresholds to be determined. The AI site is denoted consistent AI site (with respect to the PWM) if the allele with higher binding affinity also has a larger number of supporting CLIP tags (Δ*A*>0 and *B_Alt_ > B_Ref_*); otherwise, the AI site is denoted inconsistent AI site. For each PWM of an RBP, we determined the number of all consistent AI sites *N_Const_* and the number of all inconsistent AI sites *N_Inconst_*. If the PWM correctly characterized the binding specificity of the RBP, we would expect *N_Const_* > *N_Inconst_*. The excess of consistent AI sites over inconsistent AI sites was performed using a one-sided Binomial test with a null hypothesis *r = N_Const_/*(*N_Const_ + N_Inconst_*) < 0.5. We denote a PWM AI-consistent PWM if *p*<0.05 and *r*>0.5.

To determine the optimal thresholds of *α* and *δ*, we used a representative PWM (i.e., the first PWM) for each RBP and performed a grid search of parameters *α* and *δ* that maximized the number of AI-consistent PWMs. For analysis described in this paper, we used *α =* 0.8 and *δ =* 0.09 to determine AI-consistent PWMs, as the combination maximized the number of AI-consistent PWMs. With these determined thresholds, we then ranked all PWMs of each RBP based on the FDR using a single-sided binomial test (null hypothesis *r*<0.5; FDR derived from Benjamini correction for each RBP). The most significant PWM was selected as the best PWM for each RBP (Figure 5). We also used the best PWMs for each RBP to define the subset of individual consistent AI sites that most likely affect RBP binding directly (Table S4).

### Comparison of PWMs by mCross, MEME and RNAcompete

We compared the number of AI-consistent PWMs by mCross, MEME (Bailey and Elkan, 1994) and RNAcompete (Ray et al., 2013). PWMs by MEME were derived from eCLIP data for all RBPs in HepG2 and K562 cells using 100 nt sequences around CLIP tag peaks as foreground. Flanking sequences of the same size (500 nt away from peaks) were used as background. The following parameters (-dna -mod zoops -nmotifs 10 - minw 4 -maxw 7) were used to limit the motif size to 4–7 nt, allowing any number of sites per sequence. For comparison, only the first PWM was used for each RBP. In addition, we also considered 18 RBPs assayed by eCLIP that have one or more PWMs from RNAcompete for comparison with mCross. The AI-consistent PWMs were defined as described above, using the same parameters.

### Prediction of SRSF1 binding GGA clusters

We predicted clustered GGA motif sites that are bound by SRF1 using mCarts (Weyn-Vanhentenryck and Zhang, 2016; Zhang et al., 2013) to predict GGA motif clusters. Briefly, we trained the HMM using SRSF1 eCLIP peaks, extending 50nt flanking either side of every peak. Sequences without overlaps with CLIP tags were used as background. We trained one model for each cell type and identified 1,781,913 and 1,759,031 clusters for HepG2 and K562, respectively. We found substantial overlap between the two models, with 1,685,575 overlapping clusters (Figure 6C) and highly correlated cluster scores (r=0.98; Figure 6D). As such, we decided to focus on the HepG2-generated model because of its larger training set. For comparison, we also predicted conserved GGAGGA sites using branch length score (BLS) estimated from multiple alignments of 40 mammalian species (Zhang et al., 2008).

### Analysis of differential splicing upon SRSF1 knockdown

We downloaded SRSF1 shRNA knockdown and matched control data for both cell lines from ENCODE (Van Nostrand et al., 2017). RNA-seq reads were mapped with OLego (v1.1.5) using the stranded mode (Wu et al., 2013). AS quantification and differential splicing analysis upon SRSF1 knockdown were performed using Quantas as previously described (Yan et al., 2015), requiring a minimum coverage of 20 reads and FDR≤0.05. To generate RNA maps of SRSF1-dependent splicing, we used cassette exons with |ΔΨ|≥0.2 in HepG2 cells (465 exons with SRSF1-dependent inclusion and 396 exons with SRSF1-dependent exclusion). For sensitivity and positive prediction plots, we required |ΔΨ|≥0.1 to identify 373 exons with SRSF1-dependent inclusion in both HepG2 and K562 cells. Direct SRSF1 target cassette exons were predicted using overlapping GGA clusters and the exons were ranked by the cluster with the maximum motif score.

